# Claim-Level Transparency Analysis of LLM-Generated Diagnostic Reports: A Metabolic and Endocrine Biomarker Study

**DOI:** 10.64898/2026.05.03.721751

**Authors:** Andrii Yasinetsky, Caleb Geniesse, Elena Ikonomovska

## Abstract

Large language models are increasingly deployed in clinical decision-support contexts, yet systematic evaluation of their factual reliability in generating patient-specific diagnostic reports remains sparse, particularly for laboratory interpretation tasks. This study presents a controlled transparency experiment in which four frontier LLMs — Claude Sonnet 4.6, Claude Opus 4.6, GPT-5.2, and Gemini 3.1 Pro — each generated diagnostic reports for 36 patients (29 female, 7 male; aged 27–64) with biomarker profiles spanning metabolic, endocrine, and nutritional markers. A transparency engine^1^ extracted up to 50 claims per report (3,035 total), searched for supporting scientific evidence, and classified each claim as supported by science, plausible, or unsupported. Unsupported claims were uncommon: the transparency engine classified 2.7% of claims as unsupported (hereafter, the pipeline-measured hallucination rate; naive claim-level 95% Wilson CI: 2.2%–3.4%), with GPT-5.2 at the lowest observed rate (1.7%) and Claude Opus 4.6 at the highest (3.6%). However, mechanistic verification revealed a much larger plausibility gap: 915 claims (30.2%) were biologically reasonable but lacked a fully verified evidence chain, bringing the share of claims not fully supported by direct evidence to 32.9%. Gemini 3.1 Pro produced the highest plausible proportion (39.6%), suggesting a more conservative but less fully grounded reasoning profile. Although coarse support-level distributions were broadly similar across models (Cramer’s V = 0.081), claim-level analysis revealed substantial narrative divergence: 61.2% of claims were unique to a single model, and matched-claim agreement was low (Cohen’s kappa = 0.233), indicating that models generate substantively different clinical narratives for the same patient data despite comparable aggregate support profiles. These findings show that hallucination metrics alone understate the share of claims not fully verified under this protocol, and that claim-level mechanistic verification is needed to distinguish the proven from the merely plausible in metabolic and endocrine laboratory interpretation, with generalizability to other clinical domains requiring further study.

## Introduction

The integration of large language models into biomedical and clinical workflows has accelerated rapidly, with recent reviews documenting applications in diagnostics, documentation, patient communication, education, and research support [1, 2]. As these models are increasingly asked to interpret patient-specific laboratory data — particularly metabolic and endocrine biomarker panels — and recommend clinical pathways, the question of factual reliability becomes paramount. Hallucination remains a central limitation of LLMs in general [3] and in medicine specifically, where evaluations on realistic clinical tasks have identified failures in diagnostic accuracy, guideline adherence, laboratory interpretation, and robustness to misleading context [4–7]. A single unsupported claim in a diagnostic report — for instance, asserting that a nutrient deficiency causes a symptom through a mechanism not established in the literature — could mislead clinicians and harm patients.

**Figure 1.**
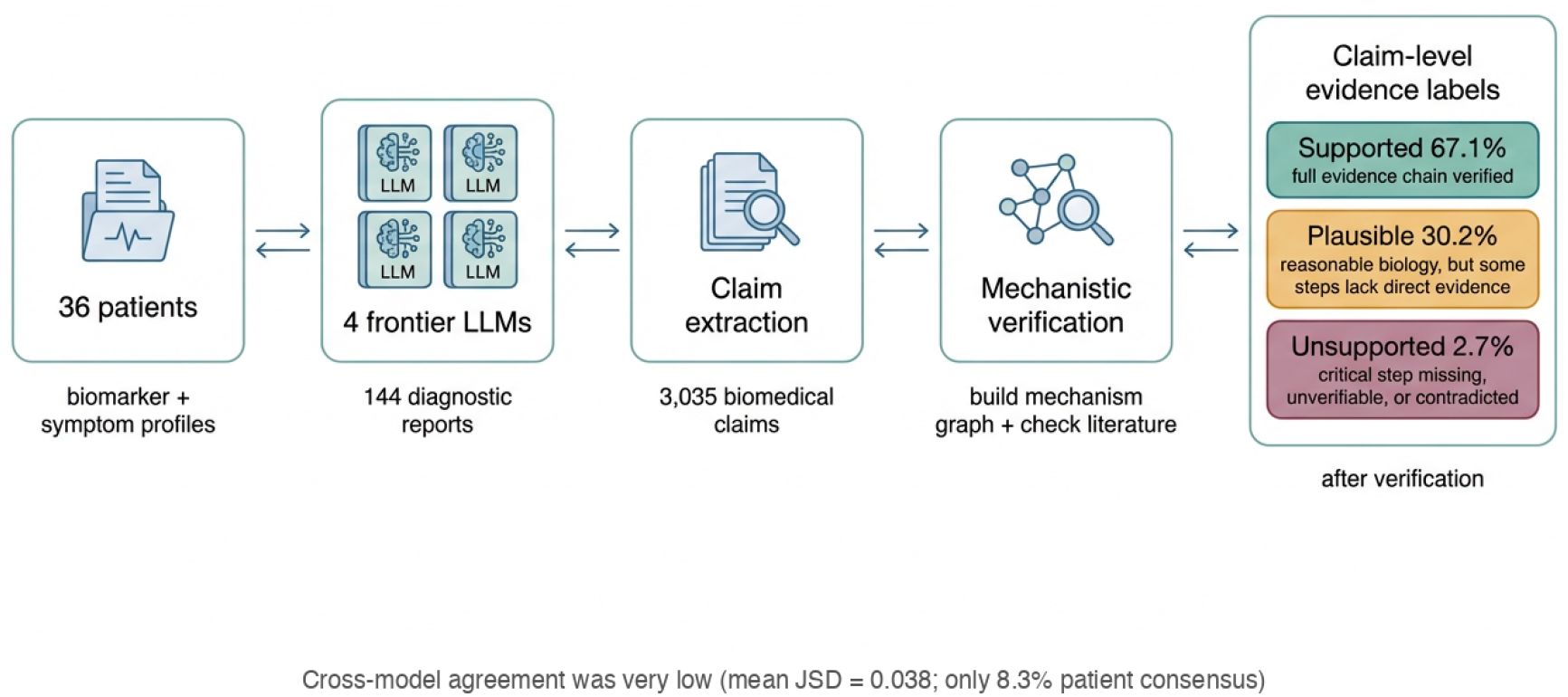
Graphical abstract illustrating the transparency experiment pipeline. Patient biomarker data from 36 individuals was processed through four frontier LLMs, producing 144 diagnostic reports. The transparency engine extracted 3,035 claims and evaluated each against the scientific literature. The overall hallucination rate was 2.7%, with variation across models ranging from 1.7% to 3.6%.

At the same time, explainability and transparency remain unresolved requirements for AI-based clinical decision support. Technical, philosophical, and ethical reviews converge on the point that aggregate performance metrics alone are insufficient when system outputs may influence patient care; context-appropriate explanations, prospective validation, and clear limits on autonomous use remain necessary [8–10].

Prior work on LLM reliability has largely emphasized benchmark performance, clinical vignettes, or summarization-oriented tasks [2–5, 11]. Emerging resources for clinical claim verification and citation integrity show that evidence-grounded evaluation is becoming tractable at the sentence and claim level [12, 13]. Comparatively little attention has been paid to the specific failure modes that arise when models generate mechanistic biomedical claims grounded in real patient data and then present those claims as integrated diagnostic narratives.

This study addresses this gap through a controlled transparency experiment. We tasked four frontier LLMs with generating diagnostic reports for 36 patients using real biomarker data, then subjected each report to automated claim extraction and evidence verification. The resulting dataset of 3,035 individually evaluated claims enables fine-grained comparison of hallucination rates, evidence quality, cross-model agreement, and the specific mechanistic pathways most prone to fabrication. The cohort of 36 patients spans an intentionally heterogeneous but still limited clinical range — including male patients and an age range of 27 to 64 years — allowing claim-level estimates of unsupported and partially supported reasoning under a single prompting protocol.

## Methods

### Patient Cohort

Thirty-six patients were selected from an existing clinical data repository. The cohort comprised 29 female and 7 male patients aged 27–64 (median 43). Biomarker panel sizes ranged from 3 to 112 analytes per patient (median approximately 63). Symptom burden varied across patients, ranging from 1 to 9 reported symptoms or clinical concerns per patient. The cohort was drawn from a metabolic and endocrine health platform; biomarker panels predominantly comprised lipid profiles, glycemic markers (HbA1c, fasting glucose), thyroid function tests, hormonal assays, and nutritional analytes (vitamin D, B12, homocysteine, ferritin). This represents a specific clinical niche rather than general medicine. All patient data was de-identified.

### Report Generation

Each patient’s data was submitted to four frontier LLMs in parallel:

- **Claude Sonnet 4.6** (Anthropic) — mid-tier reasoning model
- **Claude Opus 4.6** (Anthropic) — flagship reasoning model
- **GPT-5.2** (OpenAI, via Azure) — flagship generative model
- **Gemini 3.1 Pro** (Google) — advanced multimodal model

All models received identical prompts containing patient demographics (age, gender), symptoms, chief complaints, and relevant clinical context (e.g., diagnosed conditions, medication history, menstrual cycle timing), and a formatted biomarker list. Each model was instructed to provide root cause analysis and actionable recommendations. No system prompt was used; each model received only a user-role message. All models were invoked using their respective provider APIs with default reasoning-mode parameters: temperature, top_p, and seed were left at provider defaults and not standardized across vendors. The exact model identifiers were: `claude-sonnet-4-6` and `claude-opus-4-6` (Anthropic API), `gpt-5.2` (Azure OpenAI, endpoint API version `2025-03-01-preview`), and `gemini-3.1-pro-preview` (Google Gemini API). The maximum output token limit was set to 10,000. A single response was generated per model per patient, yielding 144 reports total (4 models × 36 patients). Reports were generated in parallel (one thread per model) during a single session in March 2026.

Report lengths varied substantially across model families. The two Anthropic models produced the longest reports (Claude Sonnet 4.6: mean 26,614 characters; Claude Opus 4.6: mean 25,958 characters), followed by GPT-5.2 (mean 12,951 characters) and Gemini 3.1 Pro (mean 8,016 characters). This three-fold difference in verbosity between the most and least verbose models is itself a meaningful variable, as longer reports present more opportunities for both supported and unsupported claims. Notably, the Anthropic models also generated the most claims per patient (Sonnet: 25.5; Opus: 26.2), compared with GPT-5.2 (18.4) and Gemini (14.2), indicating that claim density tracks with report length.

### Transparency Engine

Each of the 144 reports was processed through a transparency pipeline that performed three operations: (1) claim extraction, identifying up to 50 discrete biomedical assertions per report; (2) mechanism graph construction, mapping each claim to a directed graph of physiological nodes (biomarkers, processes, conditions, interventions) connected by mechanistic edges (causes, modulates, indicates, disrupts); and (3) evidence search, querying the scientific literature to determine whether each edge in the mechanism graph is supported.

Evidence retrieval used a web search API in academic mode, returning results from PubMed-indexed journals, preprint servers, and clinical guidelines. No date bounds were imposed on retrieval. For each claim, retrieved evidence was evaluated by an LLM acting as a medical evidence evaluator, which assigned two per-item scores: (1) a *relevance score* (0.0–1.0), reflecting how directly the evidence addresses the specific mechanism edge under evaluation (not a general quality rating); and (2) a *strength classification*: **strong** (systematic reviews, meta-analyses, well-powered RCTs, formally graded guidelines), **moderate** (RCTs or cohorts with notable limitations, consensus statements without formal grading), or **weak** (cross-sectional studies, small case series, narrative reviews, expert opinion, preprints). Up to 5 evidence items were retained per search task, sorted by relevance. Contradictory evidence was handled implicitly: if retrieved evidence contradicted a mechanistic edge, the edge was classified as unverifiable, contributing to an unsupported or plausible claim label.

Claims were classified into three support levels: *supported by science* (all mechanism paths verified), *plausible* (partial support or established biological plausibility), and *unsupported* (one or more critical mechanism paths contradicted or unverifiable). The transparency engine produced 4,864 graph components and 14,920 mechanistic edges across all claims, reflecting the depth of the verification process. All support-level classifications reported in this study are pipeline-measured: they reflect the transparency engine’s assessment based on its extraction, graph construction, and evidence retrieval capabilities, not externally validated ground-truth labels. The engine’s own accuracy introduces measurement error that is characterized in Limitations.

### Statistical Analysis

Pipeline-measured hallucination rates were computed as the proportion of claims classified as unsupported by the transparency engine, with 95% Wilson score confidence intervals [14]. These intervals treat each claim as an independent observation; because claims are nested within reports and patients and may share biomarker context, the intervals are anti-conservative and should be interpreted as naive claim-level descriptive bounds rather than as inferential estimates of true model error rates. Cross-model distributional divergence was quantified using the Jensen-Shannon divergence (JSD) [15] computed pairwise across model support-level distributions for each patient, and Cramer’s V (effect size from the model × support-level contingency table). Per-patient maximum spread was defined as the difference between the highest and lowest supported-claim proportions across the four models; a patient was considered to show model consensus when this spread was ≤15%, a threshold chosen to correspond to approximately one standard error of a binomial proportion at the median per-model claim count. Claim similarity across models was assessed using a hybrid matcher combining Jaccard index [16] on medical-term fingerprints (threshold 0.4) with cosine similarity on sentence embeddings from a pre-trained model (all-MiniLM-L6-v2; threshold 0.75). A claim pair was considered a candidate match if either criterion was met. Any claim with at least one cross-model candidate was classified as non-unique; claims with no qualifying candidate were classified as unique to their model. From the full candidate set, global 1:1 greedy matching (sorted by descending similarity) selected a non-overlapping subset of pairs for inter-model agreement analysis; each claim could contribute to at most one selected pair. Cohen’s kappa [17] was computed on these selected pairs to measure inter-model agreement on comparable clinical assertions. Patient difficulty was classified using a compound criterion combining inter-model spread and unsupported claim count: easy (0–1 models producing unsupported claims for that patient), moderate (2+ models producing unsupported claims), and hard (max spread > 40% combined with unsupported claims from *≥* 3 models).

## Results

### Overall Hallucination Rates

Across all 3,035 claims, 83 were classified as unsupported, yielding an overall hallucination rate of 2.7% (naive claim-level 95% Wilson CI: 2.2%–3.4%). The remaining claims were classified as supported by science (2,037 claims, 67.1%) or plausible (915 claims, 30.2%).

As shown in Figure 2, hallucination rates varied across models. GPT-5.2 achieved the lowest observed hallucination rate with 11 unsupported claims out of 663 (1.7%; naive 95% CI: 0.9%–2.9%), followed by Gemini 3.1 Pro with 12 out of 512 (2.3%; naive 95% CI: 1.3%–4.1%), Claude Sonnet 4.6 with 26 out of 918 (2.8%; naive 95% CI: 1.9%–4.1%), and Claude Opus 4.6 with 34 out of 942 (3.6%; naive 95% CI: 2.6%–5.0%). These intervals do not account for within-patient clustering of claims and are therefore anti-conservative; the descriptive point estimates show a two-fold spread (1.7% to 3.6%) but formal between-model inference would require clustered or multi-draw designs. Within this experiment, Opus had the highest observed unsupported-claim rate; its elevated claim count (942, the most of any model) may compound its hallucination exposure.

**Figure 2.**
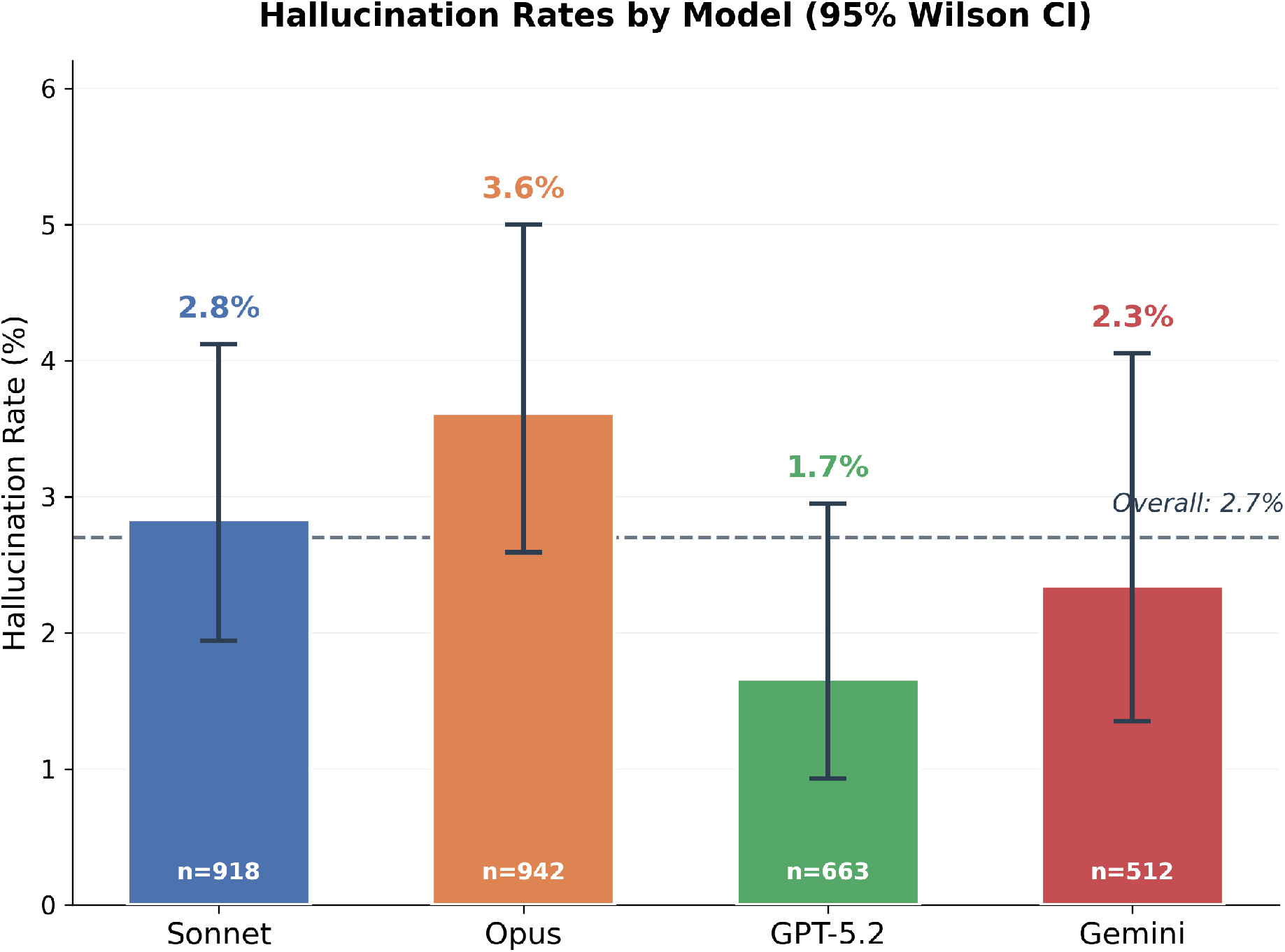
Hallucination rate by model with naive claim-level 95% Wilson intervals (not adjusted for within-patient clustering). The dashed line indicates the overall rate of 2.7%. GPT-5.2 had the lowest observed rate (1.7%), while Claude Opus 4.6 had the highest (3.6%).

### Claim Support Distribution

The distribution of support levels revealed trade-offs between models (Figure 3). GPT-5.2 generated the highest proportion of claims classified as supported by science (73.9%) while maintaining the lowest pipeline-measured unsupported rate (1.7%). Claude Opus 4.6 and Claude Sonnet 4.6 produced similar supported rates (67.3% and 67.1% respectively) and plausible rates (29.1% and 30.1%), though Opus had a higher unsupported count. Gemini 3.1 Pro produced the highest proportion of plausible claims (39.6%) and the lowest proportion of supported claims (58.0%), but maintained a relatively low unsupported rate (2.3%). This pattern suggests that Gemini may adopt a more conservative reasoning strategy, making claims that are biologically reasonable but not always directly verifiable in the literature, rather than asserting strong mechanistic links. GPT-5.2 achieved the highest pipeline-measured supported rate, while Gemini produced a higher plausible share with a comparably low unsupported rate.

**Figure 3.**
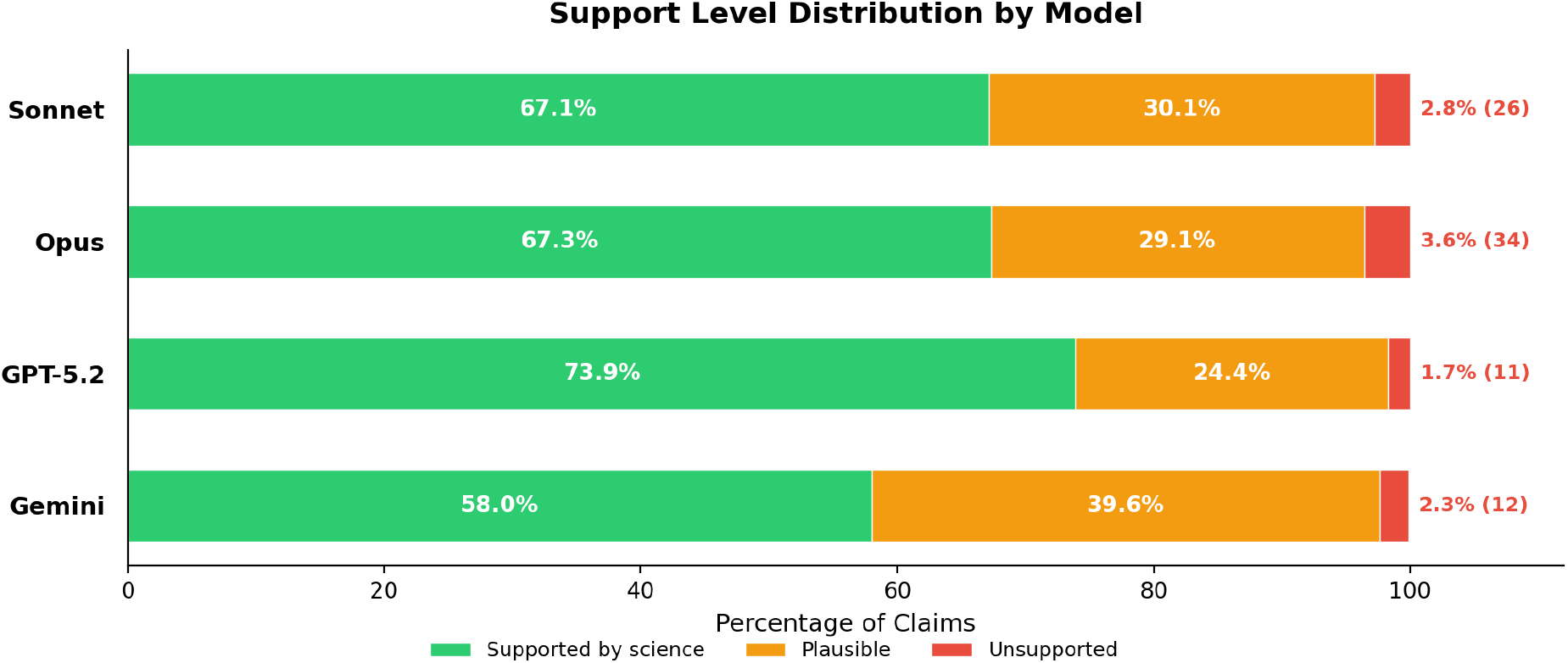
Distribution of claim support levels across models. Each bar represents the full claim set for that model. Annotations show the unsupported percentage. GPT-5.2 exhibited the highest proportion of fully supported claims (73.9%), while Gemini 3.1 Pro had the highest proportion of plausible claims (39.6%).

### The Plausible Grey Zone

The three-tier classification enabled by the transparency engine reveals a critical insight that is invisible in conventional LLM deployments. Of the 3,035 claims evaluated, 915 (30.2%) were classified as plausible — biologically reasonable assertions that have partial mechanistic support but where one or more critical pathway steps could not be fully verified against the literature. This classification is only possible because the transparency engine decomposes each claim into its constituent mechanistic edges and independently verifies each one; a claim classified as plausible might have 5 of 6 mechanism paths confirmed, with the remaining path resting on established biological plausibility rather than direct evidence.

Without mechanistic verification, these 915 plausible claims would be indistinguishable from the 2,037 fully supported claims. In a standard patient-to-AI or clinician-to-AI interaction — where a user queries a model and receives a diagnostic narrative — every claim arrives with equal confidence and no traceability. The model does not signal which assertions rest on complete evidence chains and which rely on partial mechanistic reasoning. Figure 4 illustrates this asymmetry: with the transparency engine, the claim landscape is differentiated into three actionable tiers; without it, 97.3% of claims appear uniformly “accepted,” with only the 2.7% unsupported rate as a visible risk signal.

**Figure 4.**
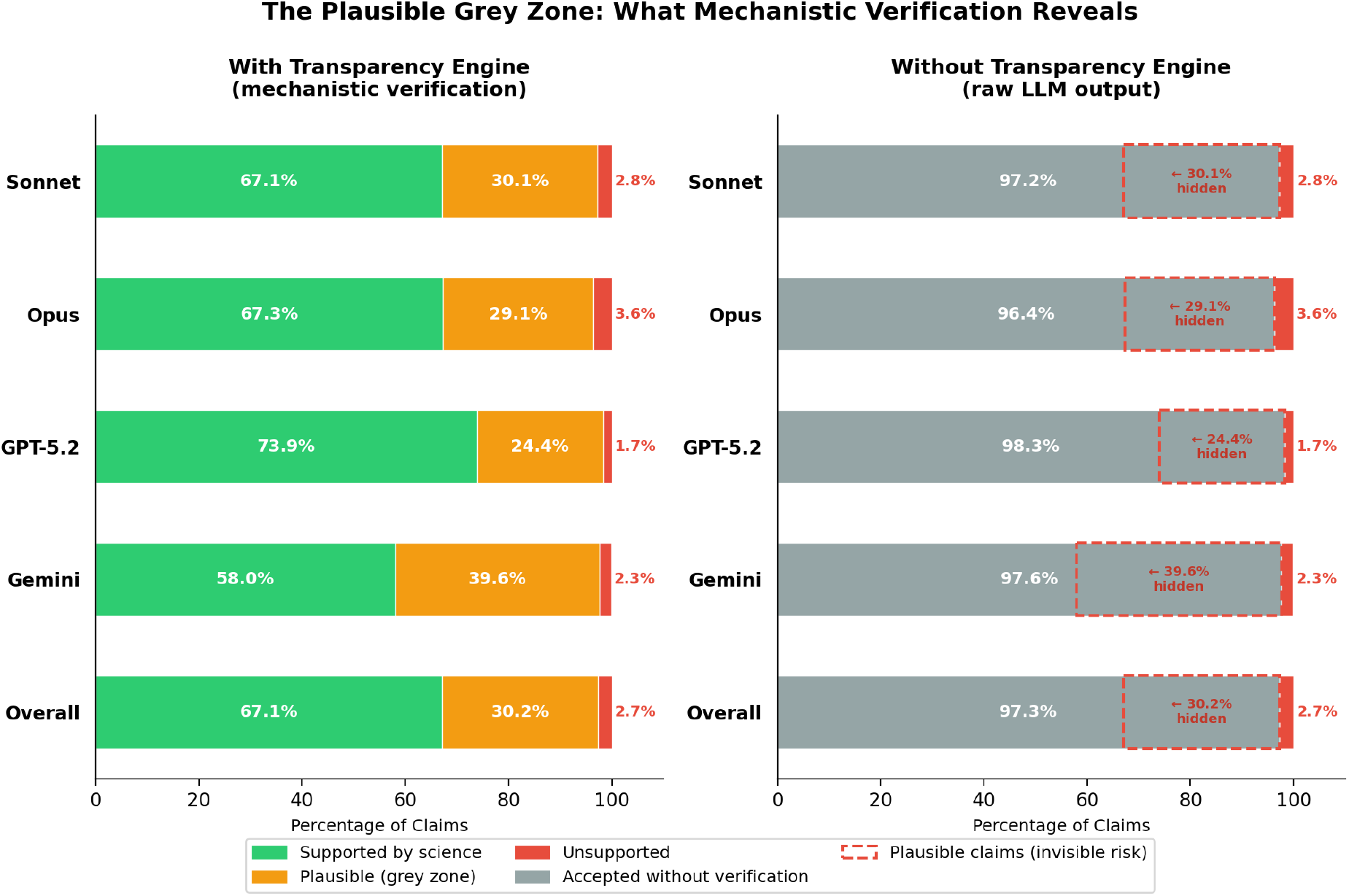
Comparison of claim visibility with and without the transparency engine. Left: mechanistic verification differentiates claims into three tiers — supported (green), plausible (orange), and unsupported (red). Right: without mechanistic decomposition, plausible claims (dashed red outline) are absorbed into the accepted category, invisible to the end user. The 30.2% plausible rate represents unverified clinical reasoning that only becomes visible through per-edge evidence evaluation.

This makes the plausible category the most clinically important grey zone in AI-assisted clinical reasoning. These are not hallucinations — they are mechanistically grounded claims that a clinician might reasonably act upon. But they carry residual uncertainty that only becomes visible when each mechanistic step is independently verified. For Gemini 3.1 Pro, which produced the highest plausible proportion (39.6%), nearly two in five claims fall into this grey zone. Even GPT-5.2, the best-performing model, generated 24.4% plausible claims — meaning that roughly one in four of its assertions, while biologically reasonable, cannot be traced to a complete evidence chain.

The clinical implication is stark: hallucination rates alone drastically understate the scope of AI reasoning that lacks a fully verified evidence chain. A reported 2.7% hallucination rate creates a false sense of safety when an additional 30.2% of claims occupy a grey zone that only mechanistic verification can illuminate.

### Evidence Quality

The transparency engine retrieved and evaluated an average of 5.0 pieces of evidence per claim across all models (Figure 5). GPT-5.2 generated claims that attracted the most evidence (5.4 per claim), while Claude Sonnet 4.6 attracted the least (4.6 per claim). Evidence relevance scores were remarkably similar across models, ranging from 0.840 (Opus) to 0.857 (GPT-5.2), suggesting that all four models generate claims that are roughly equally searchable and relevant to the existing literature.

**Figure 5.**
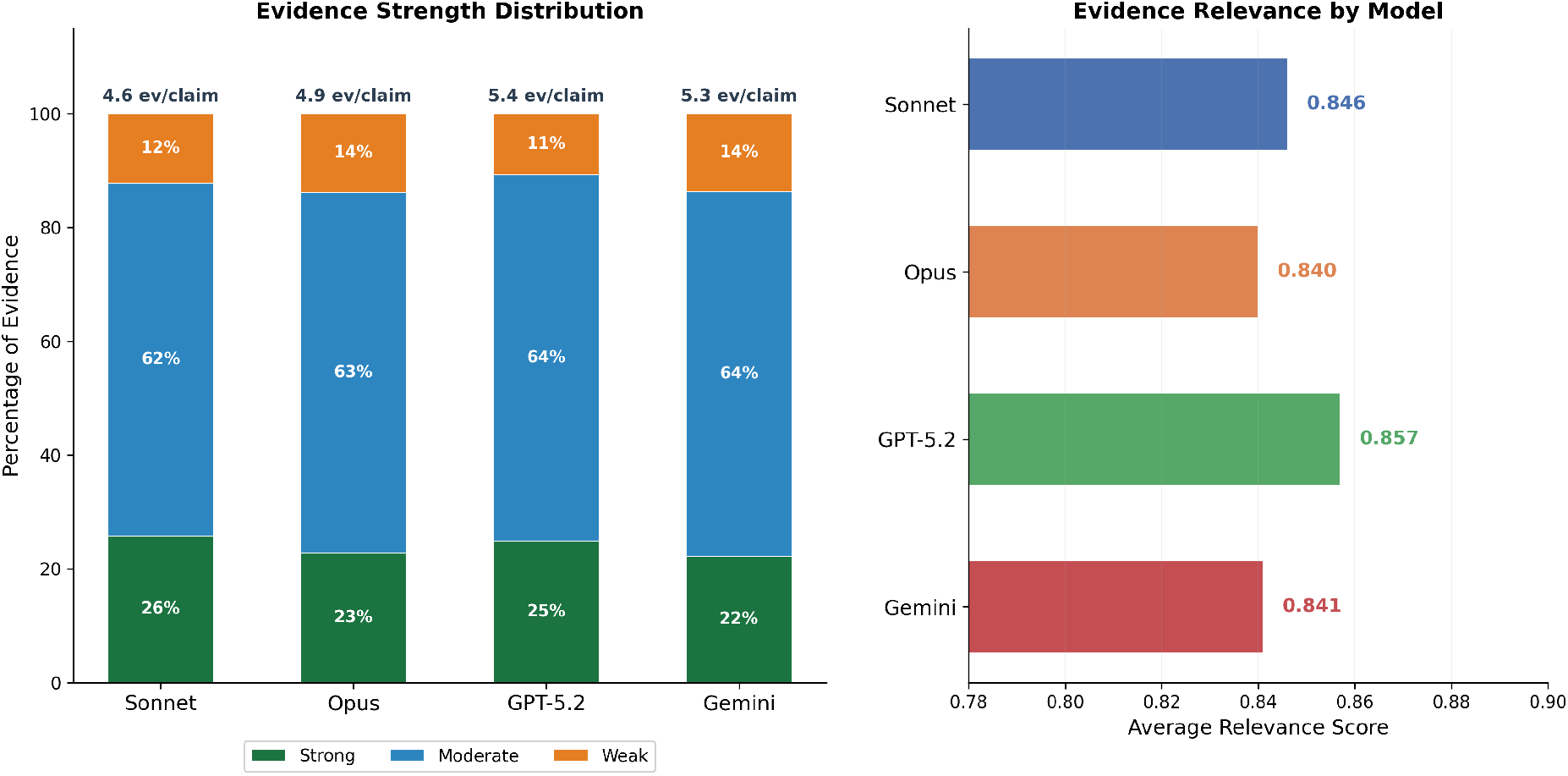
Left: Evidence quality distribution by model showing the proportion of strong, moderate, and weak evidence, with average evidence count per claim annotated above each bar. Right: Average evidence relevance scores across models.

The quality composition of evidence showed modest variation. Claude Sonnet 4.6 had the highest proportion of strong evidence (25.8%), possibly reflecting its tendency toward well-established mechanistic claims. GPT-5.2 had the lowest proportion of weak evidence (10.7%), consistent with its leading hallucination profile. Claude Opus 4.6 and Gemini 3.1 Pro showed the highest weak-evidence proportions (13.8% and 13.7% respectively), suggesting that some of their claims rest on thinner evidentiary foundations.

An analysis by support level revealed that plausible claims attracted the most evidence (5.9 per claim), compared with supported (4.6) and unsupported (4.9) claims. This may reflect the transparency engine’s more exhaustive search when evidence is mixed. The average relevance for supported claims (0.863) exceeded that for unsupported (0.819) and plausible (0.810) claims, consistent with the classification logic in which higher-relevance, higher-strength evidence contributes to a supported label. Similarly, the strong-evidence ratio for supported claims (28.5%) exceeded that for plausible (16.7%) or unsupported (17.3%) claims. These patterns reflect internal consistency of the pipeline’s scoring and classification rules rather than independent validation; external expert review would be required to assess whether the labels correspond to genuine differences in scientific grounding.

### Cross-Model Agreement

At the distributional level, models produced broadly similar support-level profiles. The mean pairwise Jensen-Shannon divergence across patient-level support distributions was 0.038, and Cramer’s V was 0.081, both indicating a negligible effect of model identity on the coarse three-category breakdown of supported, plausible, and unsupported claims. However, per-patient spread in supported-claim proportions was substantial: only 3 of 36 patients (Patients 10, 20, and 24) fell within the ≤15% maximum spread consensus threshold, yielding a patient-level agreement rate of just 8.3%.

At the claim level, narrative content diverged sharply. Among the 496 greedy-selected claim pairs (138 by both Jaccard and cosine, 345 by cosine only, 13 by Jaccard only), Cohen’s kappa was 0.233 (fair agreement), indicating that even when models discuss the same clinical topic they frequently assign different support levels. Combined with the 61.2% claim uniqueness rate (see *Claim Similarity* below), these results show that models produce substantively different diagnostic narratives despite comparable aggregate support profiles.

As Figure 6 illustrates, the degree of inter-model divergence varied substantially by patient. Patient 20, with a 2.4% spread, elicited the highest consensus: all four models produced highly similar support distributions. In contrast, Patient 17 showed 54.5% spread — the highest divergence observed — driven by Claude Sonnet 4.6 achieving 86.7% supported claims while Gemini produced only 45.5% supported. Patients 1 and 32 also showed extreme divergence (52.5% and 51.4% spread respectively). The mean maximum spread across all patients was 31.1%, indicating substantial inter-model variation as the norm rather than the exception.

**Figure 6.**
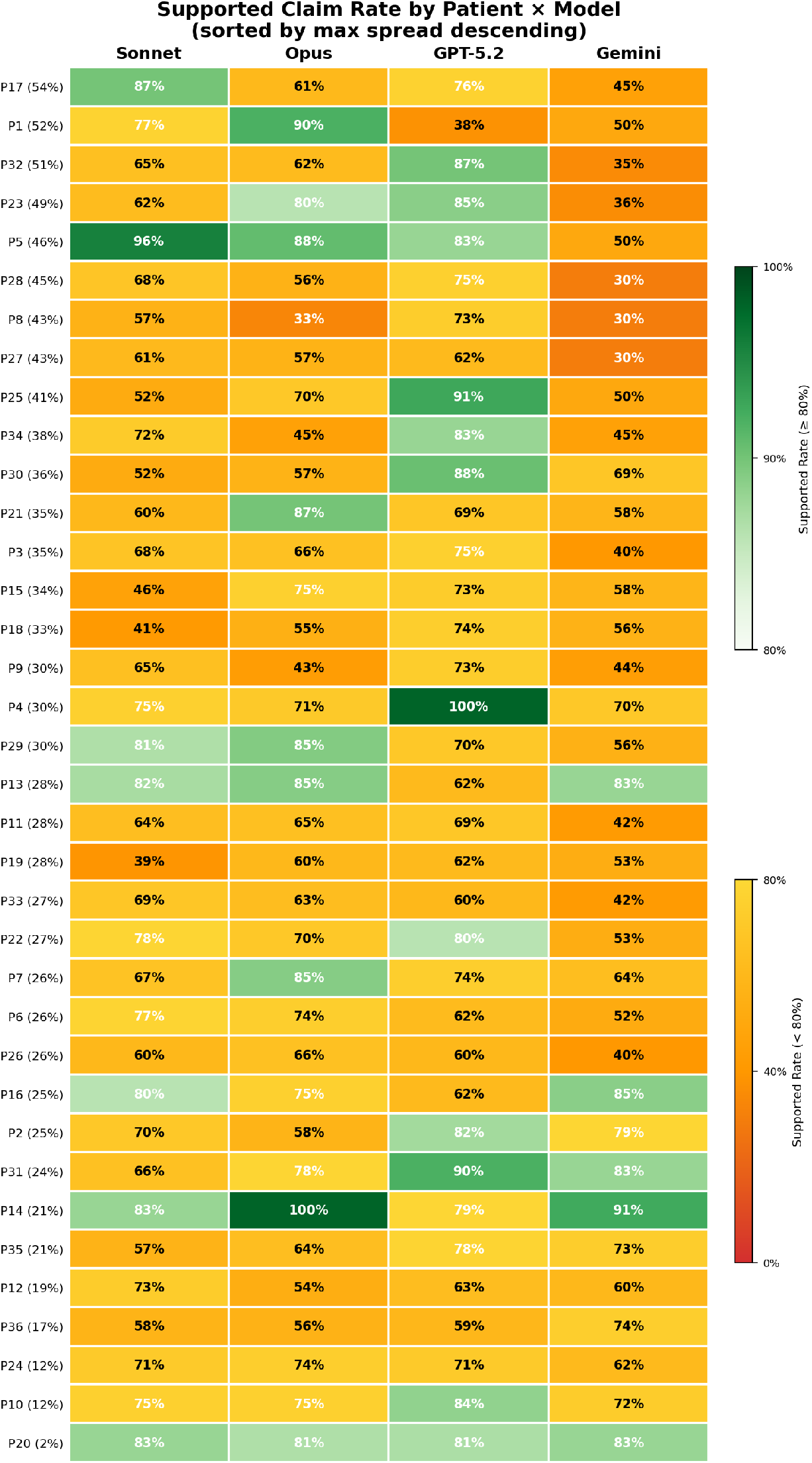
Per-patient supported claim rate and cross-model spread. The heatmap shows the proportion of supported claims for each model-patient combination, with maximum spread annotated. Patients are ordered by divergence. Twenty-eight patients were classified as easy, eight as moderate, and none as hard. The eight patients classified as moderate difficulty (Patients 2, 7, 17, 18, 21, 27, 29, and 34) tended to have higher maximum spread and in some cases unsupported claims from multiple models, suggesting that patients with higher output-derived difficulty scores also exhibited higher unsupported rates and inter-model divergence, though this association is circular since difficulty was defined using the same model outputs. No patients were classified as hard (max spread > 40% combined with unsupported claims from ≥3 models), though several came close.

### Hallucination Deep Dive

The 83 unsupported claims fell into distinct mechanistic categories. Because a single claim can contain multiple unsupported edges, the edge-level counts below sum to more than 83. The most common unsupported mechanism types were *indicates* (34 unsupported edges), *leads_to* (27), *modulates* (27), and *associated_with* (15). The most frequent unsupported node-type pairs were biomarker-to-condition (24 edges), biomarker-to-physiological_process (18), and condition-to-biomarker (8). These patterns suggest that models are most prone to hallucination when asserting that specific biomarker values indicate particular conditions or physiological processes — precisely the kind of diagnostic reasoning that is clinically consequential.

The dominance of *indicates* as the most common unsupported mechanism type is notable, as it directly mirrors the diagnostic reasoning process. The biomarker-to-condition edge type — asserting that a lab value indicates a disease state — is particularly concerning because it directly mirrors clinical diagnostic reasoning.

Among the specific unsupported claims, several thematic patterns emerged across models. Multiple unsupported claims concerned micronutrient-thyroid interactions and over-specific endocrine thresholds. Several claims about FSH thresholds for diagnosing perimenopause or menopause were marked unsupported, reflecting the models’ tendency to present rigid one-number cutoffs as definitive (e.g., “FSH > 10–12 signals perimenopause” or “FSH > 30 confirms menopause”). While elevated FSH can be consistent with menopausal transition and is used selectively in some clinical settings, consensus staging frameworks such as STRAW+10 emphasize menstrual pattern and clinical context and caution against routine reliance on single laboratory thresholds [18]; the models’ framing of these cutoffs as standalone confirmatory tests oversimplifies the diagnostic picture. Claude Opus 4.6 contributed the most unsupported claims overall (34), with errors spanning diverse domains including hormone metabolism, thyroid function, and nutrient cofactor roles. Gemini’s unsupported claims tended toward oversimplified causal assertions, such as claiming that MTHFR mutations prevent cellular B12 absorption or that TSH over 2.5 indicates thyroid sluggishness.

To illustrate the clinical stakes, four examples span the principal error types (the Addendum table lists additional cases). *Reversed causality:* Claude Sonnet 4.6 stated that “thyroid hormones suppress SHBG production” (Patient 28), reversing the established relationship — thyroid hormones are primary stimulators of hepatic SHBG synthesis via thyroid-receptor activation of HNF-4*α*, and hyperthyroidism is characteristically associated with elevated SHBG. A clinician acting on this claim could draw opposite conclusions about sex-hormone bioavailability from a patient’s thyroid status. *Compound mechanistic error:* Claude Opus 4.6 claimed that “TSH of 0.789 mIU/L rules out overt hypothyroidism as a cause of depression and elevated SHBG” (Patient 2). The TSH interpretation is correct (0.789 is euthyroid), but hypothyroidism causes *low* SHBG, not elevated; the model proposes and eliminates a physiologically impossible pathway, diverting attention from the actual causes of the patient’s elevated SHBG (4 of 8 mechanism paths broken). *Debunked physiological myth:* Claude Opus 4.6 asserted that “pregnenolone is preferentially shunted toward cortisol production at the expense of DHEA, testosterone, and progesterone under chronic stress” (Patient 33). The “pregnenolone steal” hypothesis is not supported by adrenal physiology: cortisol and DHEA are produced in separate adrenal zones (zona fasciculata and zona reticularis, respectively) with independent enzymatic regulation, and ACTH stimulation tests show concurrent rises in both steroids (2 of 3 paths broken). *Fabricated transport mechanism:* Gemini 3.1 Pro claimed that a “genetic MTHFR mutation prevents cells from absorbing B12, leaving it pooling in the blood” (Patient 9). MTHFR encodes a folate-cycle enzyme with no role in B12 cellular transport, which is mediated by transcobalamin II and ABCD4; patients with MTHFR variants actually tend toward B12 *deficiency*, not excess (2 of 4 paths broken).

### Verbosity and Hallucination

Figure 7 plots mean report length against pipeline-measured hallucination rate for the four models. Claude Opus 4.6 (25,958 characters on average) had the highest unsupported rate (3.6%), while Claude Sonnet 4.6, with the longest reports (26,614 characters), had a lower rate (2.8%). GPT-5.2’s moderate-length reports (12,951 characters) achieved the lowest rate (1.7%), and Gemini 3.1 Pro’s concise reports (8,016 characters) achieved 2.3%. With only four model-level means, no meaningful relationship between verbosity and hallucination rate can be inferred; the only safe observation is that verbosity alone does not obviously explain the observed unsupported rates. Claim count may be a confound: Opus generated 942 claims (the most) while Gemini generated only 512 (the fewest), and more claims mechanically increase the surface area for errors.

**Figure 7.**
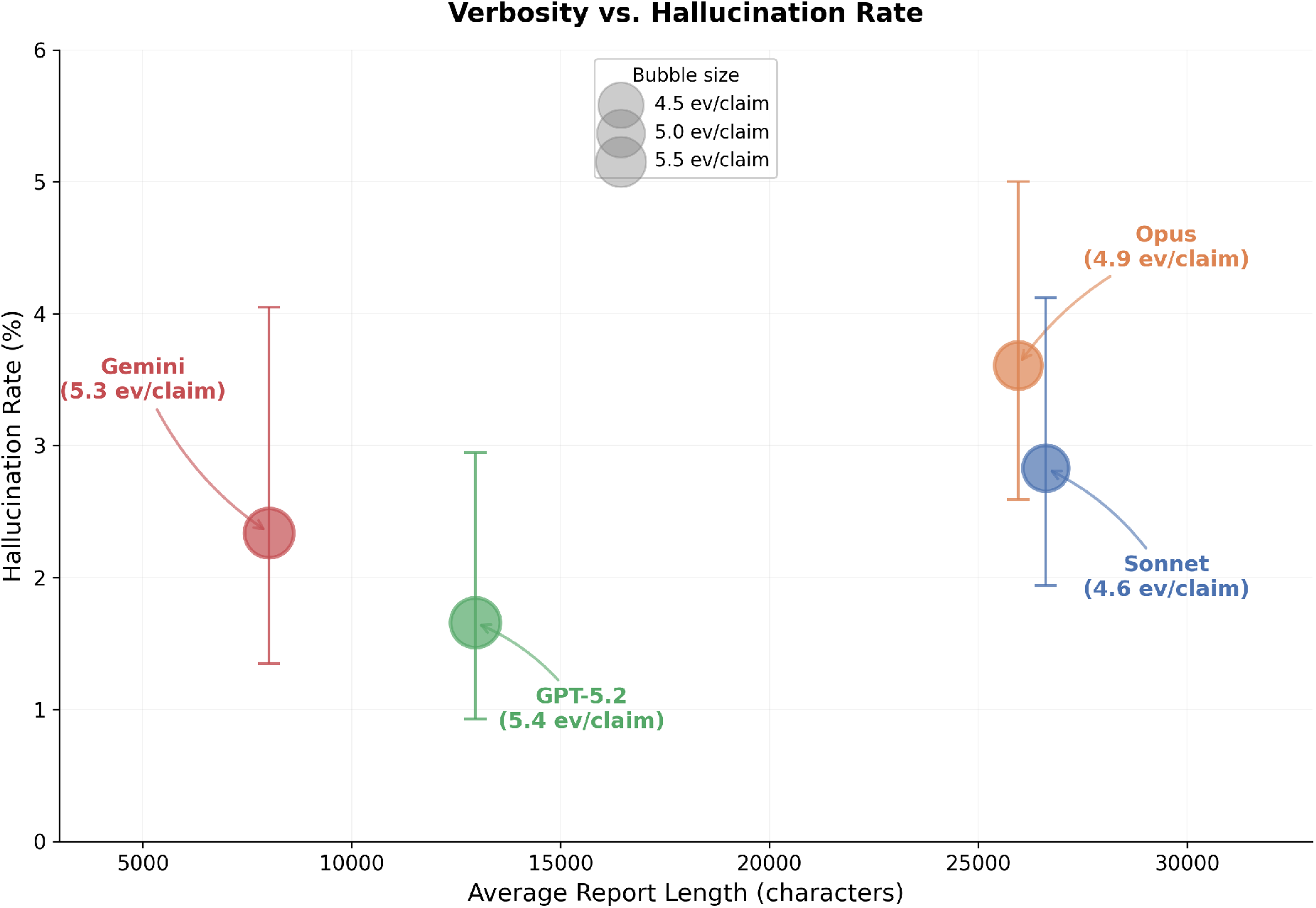
Relationship between average report length and hallucination rate. Bubble size reflects average evidence per claim. Error bars show naive claim-level 95% Wilson intervals. GPT-5.2 has the lowest observed rate at moderate verbosity, while the two Anthropic models produce similarly long reports with divergent accuracy.

### Claim Similarity

Hybrid claim matching — combining Jaccard term-overlap (threshold 0.4) with cosine similarity on sentence embeddings (threshold 0.75) — identified 1,179 claims (38.8%) with at least one cross-model candidate match and 1,856 claims (61.2%) unique to a single model. From the candidate set, global 1:1 greedy selection retained 496 non-overlapping pairs for agreement analysis, with 66.7% label agreement. Of these 496 pairs, 138 were identified by both methods, 345 by cosine similarity alone (paraphrases missed by term-overlap), and 13 by Jaccard alone. The remaining 187 non-unique claims had qualifying candidates but were excluded by greedy selection because their best candidate was already consumed by a higher-scoring pair. Per-model uniqueness rates were: Claude Sonnet 4.6: 571 (62.2%), Claude Opus 4.6: 583 (61.9%), GPT-5.2: 411 (62.0%), and Gemini 3.1 Pro: 291 (56.8%). The unsupported rate among model-unique claims was comparable to the overall rate, suggesting no dramatic risk premium for unique claims. This substantial claim diversity, combined with comparable factual accuracy, confirms that the four models are drawing on overlapping but distinct reasoning strategies when interpreting the same clinical data.

Coverage analysis revealed that matched claims were unevenly distributed across models and patients, indicating that inter-model agreement in claim substance varies widely by case.

## Discussion

This transparency experiment, encompassing 36 patients and 3,035 claims, yields four principal findings with implications for the deployment of LLMs in metabolic and endocrine laboratory interpretation.

First, the pipeline-measured hallucination rate of 2.7% across 3,035 claims is low but non-trivial in a clinical context. With the naive claim-level interval of 2.2%–3.4%, approximately 1 in 37 biomedical claims generated by frontier LLMs lacks adequate scientific support. In a typical diagnostic report containing 15–25 claims, this translates to an expected 0.4–0.7 unsupported claims per report — a frequency that, while unlikely to dominate any single report, accumulates across patient populations. The observed model-level variation (1.7% for GPT-5.2 to 3.6% for Claude Opus 4.6) represents a roughly two-fold spread in descriptive hallucination rates. Because claims are clustered within patients and each model produced only a single response per patient, these point estimates should not be over-interpreted as stable model properties; dedicated clustered or multi-draw designs would be needed to establish whether the observed spread reflects genuine between-model differences.

Second, and perhaps most consequentially, the 2.7% hallucination rate dramatically understates the broader scope of AI reasoning that lacks a fully verified evidence chain. The transparency engine’s mechanistic decomposition revealed that 30.2% of all claims — 915 out of 3,035 — occupy a plausible grey zone where biological reasoning is partially supported but at least one critical mechanistic step lacks direct evidence. Without per-edge verification, these claims are indistinguishable from fully supported ones. In any direct LLM interaction — whether a patient querying a chatbot or a clinician using an AI assistant — the model presents all claims with equal confidence. In this evaluation, the share of claims without a fully verified evidence chain was 32.9% (plausible plus unsupported), an order of magnitude larger than hallucination metrics alone suggest. This framing aligns with wider arguments that clinical AI evaluation requires more than aggregate accuracy alone and must also account for transparency and explainability in context [8–10].

Third, the agreement analysis reveals a split between distributional similarity and narrative divergence. At the aggregate level, models converge: the mean pairwise JSD of 0.038 and Cramer’s V of 0.081 indicate that the coarse split among supported, plausible, and unsupported claims is broadly comparable across models. However, at the claim level, models diverge sharply: 61.2% of claims were unique to a single model, and among the 496 greedy-selected matched pairs, label agreement was only 66.7% (Cohen’s kappa = 0.233, fair agreement). The per-patient spread in supported-claim proportions (mean 31.1%; only 3 of 36 patients within the 15% consensus threshold) further confirms that individual patients elicit markedly different reasoning from different models. Together, these findings mean that a patient receiving an AI-generated diagnostic report would encounter not just different wording but fundamentally different clinical reasoning depending on which model produced the report, even though the overall proportion of unsupported claims would be roughly similar.

Fourth, the hallucination patterns reveal a specific taxonomy of failure modes. The dominance of *indicates* edges among unsupported mechanism types (34 unsupported edges across the 83 claims) highlights a specific diagnostic failure mode: models confidently assert that biomarker values indicate conditions when the evidence is more equivocal. The concentration of errors on biomarker-to-condition and biomarker-to-physiological_process edges (42 unsupported edges) means that the most dangerous hallucinations occur precisely where clinical reasoning is most consequential — in the interpretive leap from a lab value to a diagnostic conclusion. Recurring thematic errors around micronutrient-thyroid relationships, hormonal thresholds, and nutrient-to-disease relationships suggest systematic knowledge gaps rather than random errors, which may be addressable through targeted model fine-tuning or retrieval-augmented generation [19]. The concrete examples in Section 5.6 illustrate the clinical stakes: reversed causality (thyroid hormones claimed to suppress rather than stimulate SHBG), fabricated transport mechanisms (MTHFR mutations claimed to block B12 absorption), and debunked physiological myths (“pregnenolone steal”) each represent claims that could directly alter patient management if accepted at face value. A limitation of this analysis is that all unsupported claims are weighted equally regardless of clinical severity. A speculative micronutrient cofactor mechanism and an incorrect disease-state assertion both count as one unsupported claim, but their downstream clinical consequences differ substantially. Future work should incorporate severity-weighted hallucination metrics that distinguish trivial speculative claims from assertions that could directly alter clinical management.

### Limitations

Several limitations constrain the interpretation of these results. The cohort of 36 patients remains predominantly female (30 of 36), limiting generalizability to male patients. The age range (27–64) excludes pediatric and elderly populations. Critically, the transparency engine’s own accuracy in claim extraction, graph construction, evidence retrieval, and support-level classification has not been externally validated. All rates reported in this study are pipeline-measured and may differ from rates that would be obtained by expert manual review. Claims labeled “unsupported” may in some cases reflect gaps in evidence retrieval rather than genuine fabrication, and claims labeled “supported” may include cases where the engine’s evidence matching is overly permissive. Independent validation of the pipeline against expert-annotated ground truth is a prerequisite for interpreting these rates as absolute measures of model factuality. The 50-claim-per-report extraction limit was rarely reached (average claims per report ranged from 14.2 to 26.2), but may still miss claims in the longest reports. We also evaluated one prompting template, one transparency pipeline, and one time-bounded snapshot of each vendor model; results may not generalize to other prompts, later model revisions, or alternative retrieval configurations. Finally, the inter-model comparisons are complicated by substantial differences in claim counts (512 to 942), making rate comparisons more informative than count comparisons. The cohort is drawn exclusively from a metabolic and endocrine health context; generalizability to other clinical domains (e.g., oncology, cardiology, infectious disease) remains untested. Unsupported claims are not weighted by clinical severity, meaning a speculative micronutrient mechanism and a harmful diagnostic assertion contribute equally to hallucination rates. Because each model produced only a single response per patient, the observed inter-model differences may partly reflect stochastic sampling variation rather than stable model properties. Replication with multiple draws per model, ideally with fixed seeds where supported, would be needed to establish the reliability of the reported between-model differences. Additionally, all confidence intervals reported in this study are naive claim-level Wilson intervals that treat each claim as an independent observation. In practice, claims are nested within reports and patients and share biomarker context, introducing intra-cluster correlation that makes these intervals anti-conservative. Patient-clustered bootstrap intervals or multilevel models would provide more appropriate uncertainty estimates; the intervals reported here should be read as descriptive bounds on the observed claim-level proportions, not as inferential estimates of true model error rates. Additionally, exact model snapshots may have been updated by providers between the time of generation and publication; the model identifiers and API versions reported in Methods reflect the configuration at the time of the experiment.

### Implications

For clinical deployment, these findings carry direct implications. The low but non-zero pipeline-measured unsupported rate (2.7%) supports continued human oversight of AI-generated clinical content, with particular scrutiny applied to claims that specific biomarker values indicate particular conditions — the most common failure mode identified in this study. This is consistent with prior work showing that present-day medical LLMs remain unreliable under realistic workflows or adversarial perturbation without safeguards [4–7]. More critically, the 30.2% plausible grey zone demonstrates that hallucination rate alone is a dangerously incomplete metric for clinical AI safety in this domain. Any deployment of LLMs in metabolic and endocrine diagnostic contexts without mechanistic claim verification exposes patients and clinicians to a substantial volume of biologically reasonable but incompletely evidenced reasoning — reasoning that the model itself cannot distinguish from its fully supported claims. The extreme inter-model diversity suggests that multi-model ensembling could serve as a lightweight quality signal: claims appearing in only one model’s output carry a marginally higher observed unsupported rate (2.9% vs. 2.7% overall), though the effect is small. More promisingly, retrieval-augmented generation and related grounding strategies have improved performance in biomedical LLM applications and provide a plausible route for reducing unsupported mechanistic claims [19]. Within the model set evaluated here, GPT-5.2 achieved both the highest supported-by-science rate (73.9%) and the lowest observed hallucination rate (1.7%), suggesting that model selection may influence the proportion of unsupported claims, though confirming this would require multi-draw or clustered designs.

## Conclusion

In a controlled evaluation of 3,035 claims extracted from metabolic and endocrine diagnostic reports generated by four frontier LLMs for 36 patients, the overall hallucination rate was 2.7% (naive claim-level 95% CI: 2.2%–3.4%). GPT-5.2 had the lowest observed rate (1.7%) and Claude Opus 4.6 the highest (3.6%). However, the headline hallucination rate tells only a fraction of the story. Mechanistic verification revealed that an additional 30.2% of claims — 915 out of 3,035 — occupy a plausible grey zone: biologically reasonable assertions where one or more mechanistic steps could not be fully verified against the literature. In any standard LLM interaction, these claims are presented with the same confidence as fully supported ones, meaning that 32.9% of claims in this evaluation lacked a fully verified evidence chain. Although coarse support-level distributions were broadly similar across models (Cramer’s V = 0.081), claim-level narrative content diverged sharply: 61.2% of claims were unique to a single model, matched-claim Cohen’s kappa was 0.233, and only 8.3% of patients fell within the consensus threshold. The dominant hallucination pattern involved biomarker-to-condition assertions, particularly around thyroid function, hormonal thresholds, and nutrient cofactor roles. These results demonstrate that hallucination rates alone are an insufficient safety metric for clinical AI in this domain, and further work is needed to weight claims by clinical severity. Without mechanistic decomposition that independently verifies each step in a claim’s reasoning chain, a large share of clinically relevant uncertainty remains invisible in ordinary patient-to-AI and clinician-to-AI interaction. LLM-generated diagnostic reports require not just human oversight, but systematic mechanistic verification to distinguish the proven from the merely plausible.

## Ethics Statement

The data underlying this study are derived from real-world patient encounters and contain protected health information. Due to privacy and contractual restrictions, the underlying transcripts and associated encounter-level data cannot be shared publicly or on request. All data were anonymized (scrubbed of any patient-revealing information).

## Data Availability

The per-claim support-level classifications, cross-model agreement statistics, and aggregate results reported in this paper are available in the supplementary materials. The underlying patient-level biomarker data and generated reports cannot be released due to the privacy restrictions described above.

## Code Availability

The system evaluated in this study is a proprietary platform and the underlying code is not publicly available.

## Addendum

### Prompt Template

All four models received the same prompt for each patient, constructed from the patient’s demographics, relevant clinical context (e.g., diagnosed conditions, medication history, menstrual-cycle timing), symptoms, and biomarker panel. No system prompt was used — the models received only the user message. The template is reproduced below, with placeholders shown in braces:

~~~
I’m a {age}-year-old {gender}. {patient_context} For the past few months I’ve been dealing with
<→ {symptoms}. It’s been really affecting my daily life and I want to get to the bottom of
<→ what’s going on.
I went to my doctor and got a comprehensive blood panel done. Here are all my results:
{biomarker list, one per line: “-Name: value unit”}
I’d really like a thorough analysis of what might be going on with my health. Based on these lab
<→ results and my symptoms, can you explain:
- What the possible root causes of my symptoms could be
- How the different abnormal biomarkers might be connected to each other
- What underlying mechanisms or conditions could explain the pattern I’m seeing
- Which findings are most concerning and why
-Provide specific recommendations and next steps for me to take to address the root causes Please give me your full analysis based on what I’ve shared. Don’t ask me any follow-up
<→ questions --just give me your best assessment with as much detail as you can.
~~~

### Patient Profiles and Model Focus Areas

The following summaries describe the clinical input provided to each model (demographics, biomarker panel, symptoms and clinical context) and the dominant analytical themes that appeared in the resulting reports. All patient data was de-identified prior to analysis.

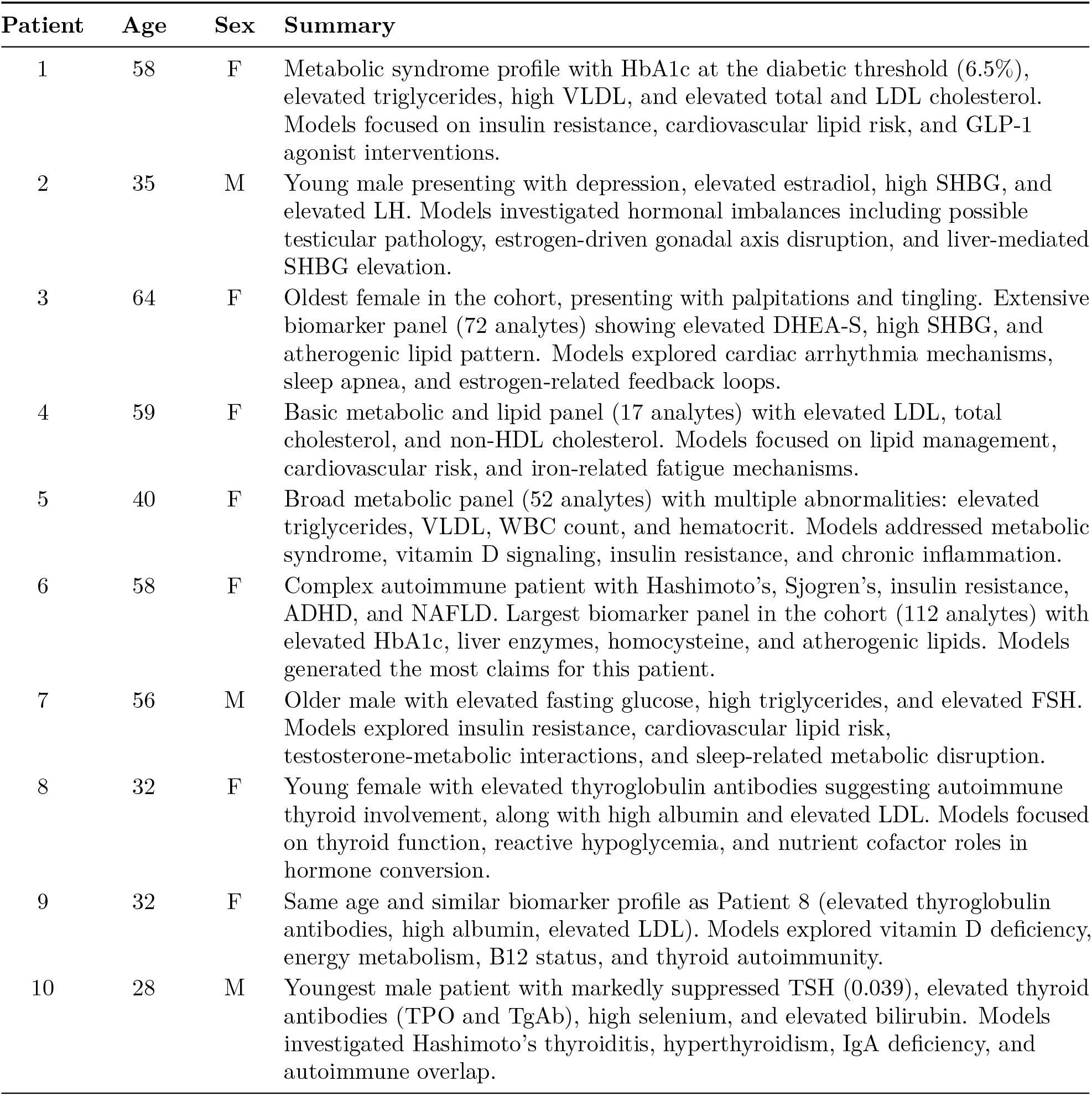
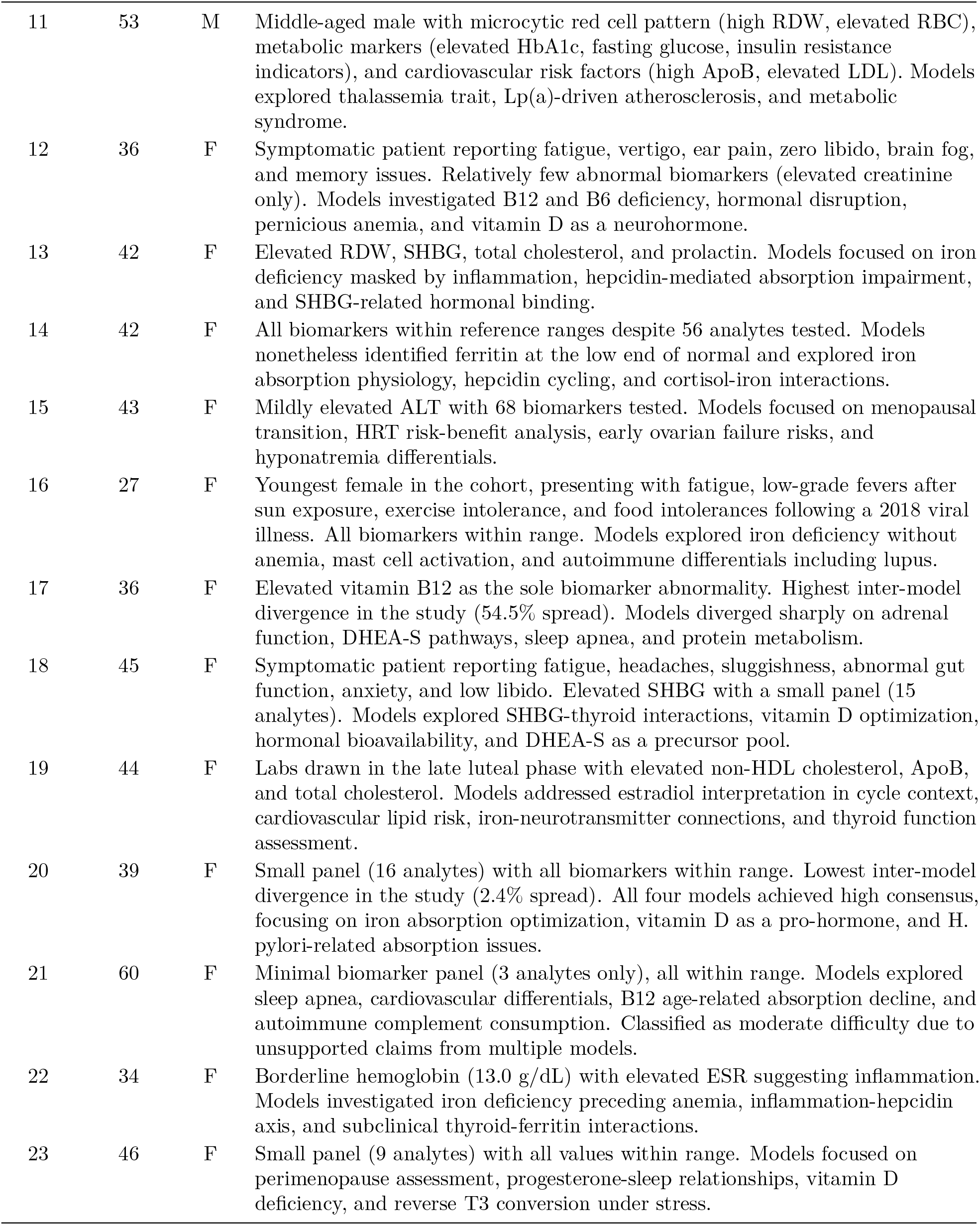
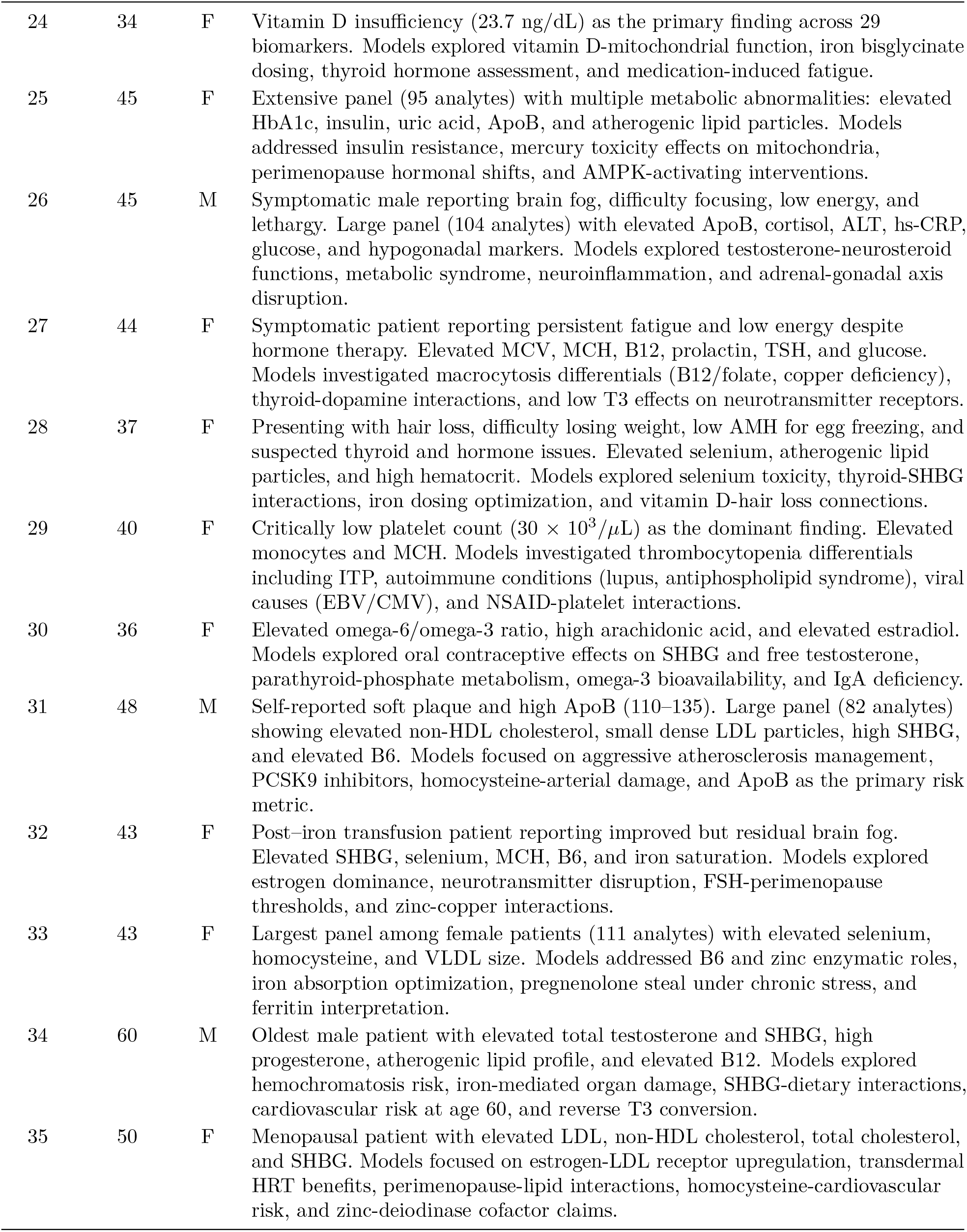
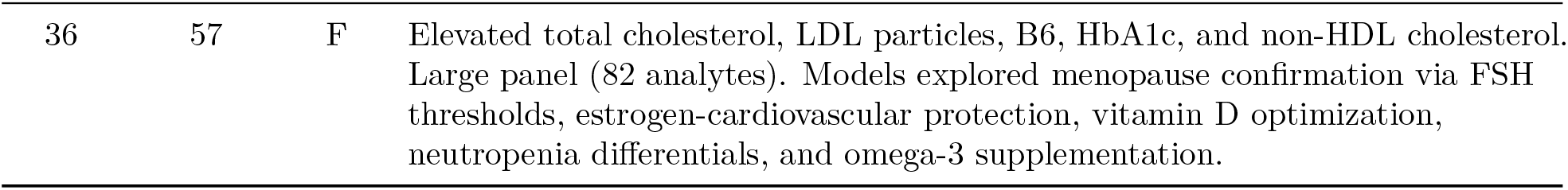

| Patient | Age | Sex | Summary |
| --- | --- | --- | --- |
| 1 | 58 | F | Metabolic syndrome profile with HbA1c at the diabetic threshold (6.5%), elevated triglycerides, high VLDL, and elevated total and LDL cholesterol. Models focused on insulin resistance, cardiovascular lipid risk, and GLP-1 agonist interventions. |
| 2 | 35 | M | Young male presenting with depression, elevated estradiol, high SHBG, and elevated LH. Models investigated hormonal imbalances including possible testicular pathology, estrogen-driven gonadal axis disruption, and liver-mediated SHBG elevation. |
| 3 | 64 | F | Oldest female in the cohort, presenting with palpitations and tingling. Extensive biomarker panel (72 analytes) showing elevated DHEA-S, high SHBG, and atherogenic lipid pattern. Models explored cardiac arrhythmia mechanisms, sleep apnea, and estrogen-related feedback loops. |
| 4 | 59 | F | Basic metabolic and lipid panel (17 analytes) with elevated LDL, total cholesterol, and non-HDL cholesterol. Models focused on lipid management, cardiovascular risk, and iron-related fatigue mechanisms. |
| 5 | 40 | F | Broad metabolic panel (52 analytes) with multiple abnormalities: elevated triglycerides, VLDL, WBC count, and hematocrit. Models addressed metabolic syndrome, vitamin D signaling, insulin resistance, and chronic inflammation. |
| 6 | 58 | F | Complex autoimmune patient with Hashimoto's, Sjogren's, insulin resistance, ADHD, and NAFLD. Largest biomarker panel in the cohort (112 analytes) with elevated HbA1c, liver enzymes, homocysteine, and atherogenic lipids. Models generated the most claims for this patient. |
| 7 | 56 | M | Older male with elevated fasting glucose, high triglycerides, and elevated FSH. Models explored insulin resistance, cardiovascular lipid risk, testosterone-metabolic interactions, and sleep-related metabolic disruption. |
| 8 | 32 | F | Young female with elevated thyroglobulin antibodies suggesting autoimmune thyroid involvement, along with high albumin and elevated LDL. Models focused on thyroid function, reactive hypoglycemia, and nutrient cofactor roles in hormone conversion. |
| 9 | 32 | F | Same age and similar biomarker profile as Patient 8 (elevated thyroglobulin antibodies, high albumin, elevated LDL). Models explored vitamin D deficiency, energy metabolism, B12 status, and thyroid autoimmunity. |
| 10 | 28 | M | Youngest male patient with markedly suppressed TSH (0.039), elevated thyroid antibodies (TPO and TgAb), high selenium, and elevated bilirubin. Models investigated Hashimoto's thyroiditis, hyperthyroidism, IgA deficiency, and autoimmune overlap. |
| 11 | 53 | M | Middle-aged male with microcytic red cell pattern (high RDW, elevated RBC), metabolic markers (elevated HbA1c, fasting glucose, insulin resistance indicators), and cardiovascular risk factors (high ApoB, elevated LDL). Models explored thalassemia trait, Lp(a)-driven atherosclerosis, and metabolic syndrome. |
| 12 | 36 | F | Symptomatic patient reporting fatigue, vertigo, ear pain, zero libido, brain fog, and memory issues. Relatively few abnormal biomarkers (elevated creatinine only). Models investigated B12 and B6 deficiency, hormonal disruption, pernicious anemia, and vitamin D as a neurohormone. |
| 13 | 42 | F | Elevated RDW, SHBG, total cholesterol, and prolactin. Models focused on iron deficiency masked by inflammation, hepcidin-mediated absorption impairment, and SHBG-related hormonal binding. |
| 14 | 42 | F | All biomarkers within reference ranges despite 56 analytes tested. Models nonetheless identified ferritin at the low end of normal and explored iron absorption physiology, hepcidin cycling, and cortisol-iron interactions. |
| 15 | 43 | F | Mildly elevated ALT with 68 biomarkers tested. Models focused on menopausal transition, HRT risk-benefit analysis, early ovarian failure risks, and hyponatremia differentials. |
| 16 | 27 | F | Youngest female in the cohort, presenting with fatigue, low-grade fevers after sun exposure, exercise intolerance, and food intolerances following a 2018 viral illness. All biomarkers within range. Models explored iron deficiency without anemia, mast cell activation, and autoimmune differentials including lupus. |
| 17 | 36 | F | Elevated vitamin B12 as the sole biomarker abnormality. Highest inter-model divergence in the study (54.5% spread). Models diverged sharply on adrenal function, DHEA-S pathways, sleep apnea, and protein metabolism. |
| 18 | 45 | F | Symptomatic patient reporting fatigue, headaches, sluggishness, abnormal gut function, anxiety, and low libido. Elevated SHBG with a small panel (15 analytes). Models explored SHBG-thyroid interactions, vitamin D optimization, hormonal bioavailability, and DHEA-S as a precursor pool. |
| 19 | 44 | F | Labs drawn in the late luteal phase with elevated non-HDL cholesterol, ApoB, and total cholesterol. Models addressed estradiol interpretation in cycle context, cardiovascular lipid risk, iron-neurotransmitter connections, and thyroid function assessment. |
| 20 | 39 | F | Small panel (16 analytes) with all biomarkers within range. Lowest inter-model divergence in the study (2.4% spread). All four models achieved high consensus, focusing on iron absorption optimization, vitamin D as a pro-hormone, and H. pylori-related absorption issues. |
| 21 | 60 | F | Minimal biomarker panel (3 analytes only), all within range. Models explored sleep apnea, cardiovascular differentials, B12 age-related absorption decline, and autoimmune complement consumption. Classified as moderate difficulty due to unsupported claims from multiple models. |
| 22 | 34 | F | Borderline hemoglobin (13.0 g/dL) with elevated ESR suggesting inflammation. Models investigated iron deficiency preceding anemia, inflammation-hepcidin axis, and subclinical thyroid-ferritin interactions. |
| 23 | 46 | F | Small panel (9 analytes) with all values within range. Models focused on perimenopause assessment, progesterone-sleep relationships, vitamin D deficiency, and reverse T3 conversion under stress. |
| 24 | 34 | F | Vitamin D insufficiency (23.7 ng/dL) as the primary finding across 29 biomarkers. Models explored vitamin D-mitochondrial function, iron bisglycinate dosing, thyroid hormone assessment, and medication-induced fatigue. |
| 25 | 45 | F | Extensive panel (95 analytes) with multiple metabolic abnormalities: elevated HbA1c, insulin, uric acid, ApoB, and atherogenic lipid particles. Models addressed insulin resistance, mercury toxicity effects on mitochondria, perimenopause hormonal shifts, and AMPK-activating interventions. |
| 26 | 45 | M | Symptomatic male reporting brain fog, difficulty focusing, low energy, and lethargy. Large panel (104 analytes) with elevated ApoB, cortisol, ALT, hs-CRP, glucose, and hypogonadal markers. Models explored testosterone-neurosteroid functions, metabolic syndrome, neuroinflammation, and adrenal-gonadal axis disruption. |
| 27 | 44 | F | Symptomatic patient reporting persistent fatigue and low energy despite hormone therapy. Elevated MCV, MCH, B12, prolactin, TSH, and glucose. Models investigated macrocytosis differentials (B12/folate, copper deficiency), thyroid-dopamine interactions, and low T3 effects on neurotransmitter receptors. |
| 28 | 37 | F | Presenting with hair loss, difficulty losing weight, low AMH for egg freezing, and suspected thyroid and hormone issues. Elevated selenium, atherogenic lipid particles, and high hematocrit. Models explored selenium toxicity, thyroid-SHBG interactions, iron dosing optimization, and vitamin D-hair loss connections. |
| 29 | 40 | F | Critically low platelet count ($30 \times 10^3/\mu\text{L}$) as the dominant finding. Elevated monocytes and MCH. Models investigated thrombocytopenia differentials including ITP, autoimmune conditions (lupus, antiphospholipid syndrome), viral causes (EBV/CMV), and NSAID-platelet interactions. |
| 30 | 36 | F | Elevated omega-6/omega-3 ratio, high arachidonic acid, and elevated estradiol. Models explored oral contraceptive effects on SHBG and free testosterone, parathyroid-phosphate metabolism, omega-3 bioavailability, and IgA deficiency. |
| 31 | 48 | M | Self-reported soft plaque and high ApoB (110–135). Large panel (82 analytes) showing elevated non-HDL cholesterol, small dense LDL particles, high SHBG, and elevated B6. Models focused on aggressive atherosclerosis management, PCSK9 inhibitors, homocysteine-arterial damage, and ApoB as the primary risk metric. |
| 32 | 43 | F | Post-iron transfusion patient reporting improved but residual brain fog. Elevated SHBG, selenium, MCH, B6, and iron saturation. Models explored estrogen dominance, neurotransmitter disruption, FSH-perimenopause thresholds, and zinc-copper interactions. |
| 33 | 43 | F | Largest panel among female patients (111 analytes) with elevated selenium, homocysteine, and VLDL size. Models addressed B6 and zinc enzymatic roles, iron absorption optimization, pregnenolone steal under chronic stress, and ferritin interpretation. |
| 34 | 60 | M | Oldest male patient with elevated total testosterone and SHBG, high progesterone, atherogenic lipid profile, and elevated B12. Models explored hemochromatosis risk, iron-mediated organ damage, SHBG-dietary interactions, cardiovascular risk at age 60, and reverse T3 conversion. |
| 35 | 50 | F | Menopausal patient with elevated LDL, non-HDL cholesterol, total cholesterol, and SHBG. Models focused on estrogen-LDL receptor upregulation, transdermal HRT benefits, perimenopause-lipid interactions, homocysteine-cardiovascular risk, and zinc-deiodinase cofactor claims. |
| 36 | 57 | F | Elevated total cholesterol, LDL particles, B6, HbA1c, and non-HDL cholesterol. Large panel (82 analytes). Models explored menopause confirmation via FSH thresholds, estrogen-cardiovascular protection, vitamin D optimization, neutropenia differentials, and omega-3 supplementation. |

### Clinically Consequential Unsupported Claims

The following table presents seven unsupported claims selected from the 83 total for their potential to alter clinical management if trusted without verification. Claims were chosen to span the principal error types observed: reversed causality, fabricated mechanisms, debunked physiological myths, and false diagnostic thresholds.

| Model | Pt | Verbatim Claim | Transparency Engine Finding | Clinical Risk |
| --- | --- | --- | --- | --- |
| Sonnet | 28 | “Thyroid hormones suppress SHBG production” | Reversed causality. Thyroid hormones stimulate hepatic SHBG synthesis via TR/HNF-4α; hyperthyroidism → high SHBG, hypothyroidism → low SHBG. 1/3 paths broken. | Opposite conclusions about sex-hormone bioavailability from thyroid status. |
| Opus | 2 | “TSH of 0.789 mIU/L rules out overt hypothyroidism as a cause of depression and elevated SHBG” | TSH interpretation correct (euthyroid), but hypothyroidism causes *low* SHBG, not elevated. Model proposes and eliminates a physiologically impossible pathway. 4/8 paths broken. | Incomplete differential diagnosis; actual causes of elevated SHBG overlooked. |
| Opus | 33 | “Pregnenolone is preferentially shunted toward cortisol production at the expense of DHEA, testosterone, and progesterone under chronic stress” | “Pregnenolone steal” is unsupported. Cortisol (zona fasciculata) and DHEA (zona reticularis) are produced in separate adrenal zones with independent enzymatic regulation. 2/3 paths broken. | Supplementation strategies targeting a non-existent mechanism; actual HPA/HPG axis dysregulation missed. |
| Opus | 2 | “Cannabis use is associated with elevated estrogen and suppressed testosterone” | THC suppresses the HPG axis, leading to *reduced* estrogen (not elevated). Testosterone suppression is plausible. 1/4 paths broken. | Wrong directional guidance on hormonal effects of cannabis use. |
| Gemini | 1 | “Optimal TSH is between 0.5 and 2.5” | No evidence that narrowing the standard 0.4–4.5 range improves outcomes. Large cohort studies (>160k patients) show no cardiovascular benefit; in older adults, TSH up to 7.0 may be protective. 2/4 paths broken. | Unnecessary levothyroxine in euthyroid patients, risking iatrogenic hyperthyroidism. |
| Sonnet | 17 | “DHEA-S is the precursor to both cortisol and sex hormones” | DHEA-S is a precursor to sex hormones but not cortisol. They are parallel products diverging from pregnenolone (Δ5 vs Δ4 pathways). 2/4 paths broken. | Attempts to manipulate DHEA-S to affect cortisol, which is physiologically impossible. |
| Gemini | 9 | “Genetic MTHFR mutation prevents cells from absorbing B12, leaving it pooling in the blood” | MTHFR encodes a folate-cycle enzyme with no role in B12 transport (mediated by TCN2/ABCD4). MTHFR carriers tend toward B12 *deficiency*, not excess. 2/4 paths broken. | Avoidance of needed B12 supplementation; unnecessary workup for non-existent high B12. |

## Footnotes

1 The transparency engine is a proprietary mechanistic reasoning system developed by Diadia. It decomposes biomedical claims into directed graphs of physiological mechanisms, independently verifies each mechanistic step against the scientific literature, and derives support classifications through deterministic graph analysis. The engine underpins every root cause analysis in Diadia’s clinical platform, providing full traceability from each diagnostic conclusion back to the specific evidence that supports or contradicts it.

